# Prion-like spreading of Alzheimer’s disease within the brain’s connectome

**DOI:** 10.1101/529438

**Authors:** Sveva Fornari, Amelie Schäfer, Mathias Jucker, Alain Goriely, Ellen Kuhl

## Abstract

The prion hypothesis states that misfolded proteins can act as infectious agents that trigger the misfolding and aggregation of healthy proteins to transmit a variety of neurodegenerative diseases. Increasing evidence suggests that pathogenic proteins in Alzheimer’s disease adapt prion-like mechanisms and spread across the brain along an anatomically connected network. Local kinetics models of protein misfolding and global network models of protein diffusion provide valuable insight into the dynamics of prion-like diseases. Yet, to date, these models have not been combined to simulate how pathological proteins multiply and spread across the human brain. Here we model the prion-like spreading of Alzheimer’s disease by combining misfolding kinetics and network diffusion through a connectivity-weighted Laplacian graph created from 418 brains of the Human Connectome Project. The nodes of the graph represent anatomic regions of interest and the edges represent their connectivity, weighted by the mean fiber number divided by the mean fiber length. We show that our brain network model correctly predicts the neuropathological pattern of Alzheimer’s disease and captures the key characteristic features of whole brain models at a fraction of their computational cost. To illustrate the potential of brain network modeling in neurodegeneration, we simulate biomarker curves, infection times, and two promising therapeutic strategies to delay the onset of neurodegeneration: reduced production and increased clearance of misfolded protein.

PACS numbers: 87.10.Ed, 87.15.hj, 87.16.Ac, 87.19.L-, 87.19.lp, 87.19.xr

## Introduction

A major advance in our understanding of the brain has been the realization that the brain is organized as a network, both at the physical and functional levels [1]. This quiet revolution has been made possible by the parallel development of medical imagng and network theory [2]. Methods originating from graph theory are now routinely used to study various aspects of brain function and the prevalent dogma is that the brain operates as an efficient, modular, dynamic network with strongly connected hubs [3]. This network is optimized to quickly transmit electrical signals, but, unfortunately, also toxic molecules that rapidly spread within the brain’s connectome [4]. Studies have shown that the eigenmodes of the brain network’s graph Laplacian are correlated to atrophy in Alzheimer’s disease [4], and probabilistic epidemiological models have used the network to explain transference mechanisms [5].

The current prevalent theory for neurodegenerative disorders is the prion-like paradigm [6] in which degeneration is caused by the invasion and conformational autocatalytic conversion of misfolded proteins [7]. In Alzheimer’s disease, tau protein is believed to act in a prion-like manner [8]: it misfolds and becomes a toxic template on which healthy proteins misfold, become toxic themselves, and grow into increasingly larger aggregates [9]. Tau is an intracellular protein that primarily spreads along axonal pathways [10]. This creates a remarkably consistent and predictable pattern [11]. Figure 1, top, illustrates the typical spatio-temporal pattern of tau protein misfolding in Alzheimer’s disease inferred from neuropathological observations of hundreds of human brains [12].

**FIG. 1:**
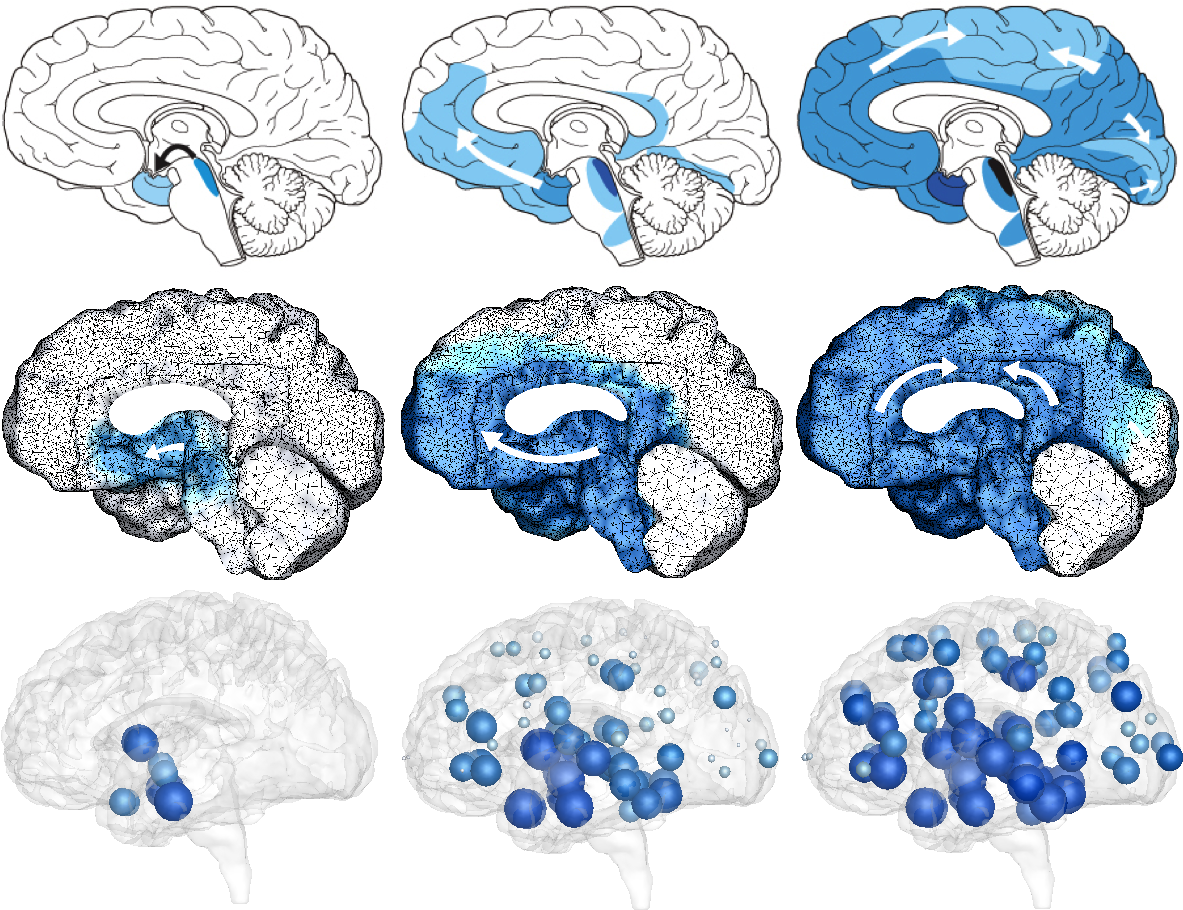
Typical pattern of tau protein misfolding in Alzheimer’s disease. Misfolded tau proteins occur first in the locus coeruleus and transentorhinal layer from where they spread to the transentorhinal region and the proper entorhinal cortex and ultimately affect all interconnected neocortical brain regions. Neuropathological observation (top) [12], continuum model (middle) [13], and network model (bottom).

Throughout the past decade, three conceptually different models have emerged to simulate the physics of protein misfolding and transport: *(i)* kinetic growth and fragmentation models to study the interaction of aggregates of different sizes using a set of ordinary differential equations [14]; *(ii)* network diffusion models to study the prion-like spreading of misfolded proteins using graph theories [4]; and *(iii)* reaction-diffusion continuum models to study the spatio-temporal evolution of pathogenic proteins using partial differential equations [15]. Figure 1, middle, shows that continuum models with nonlinear reaction and anisotropic diffusion accurately predict the typical pattern of tau protein misfolding in Alzheimer’s disease [13]. This simulation used a Fisher-Kolmogorov type equation [16, 17], discretized with 400,000 tetrahedral finite elements and 80,000 degrees of freedom. While the continuum model displays an excellent agreement with neuropathological observations, it is computationally expensive and impractical for the quick assessment of a variety of disease scenarios. The objective of this study is therefore to create an efficient and robust simulation tool that captures the key characteristic features of pathogenic proteins in Alzheimer’s disease by combining misfolding kinetics and network diffusion through a connectivity-weighted graph from the Human Connectome Project. Figure 1, bottom, shows that–even with three orders of magnitude fewer degrees of freedom– this dynamic network model accurately predicts the typical spatio-temporal pattern of tau protein misfolding.

### Kinetic model

To model the misfolding of tau protein, we consider the simplest possible kinetic model that accounts for two protein configurations, the natural healthy state *p* and the misfolded state 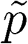 [18, 19]. In this model, misfolded proteins recruit healthy proteins at a rate *k*_11_′, healthy proteins bind to misfolded proteins and adopt their conformation at a rate *k*_1_′ _2_′, and the resulting polymer fragments into infectious seeds at a rate *k*_2_′ _2_,

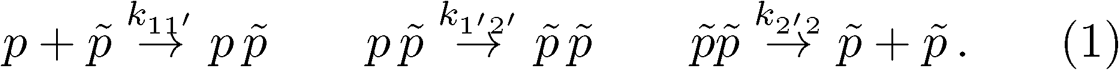

For simplicity, we collectively represent the conformational conversion from the healthy to the misfolded state as a single step through the rate constant *k*_12_,

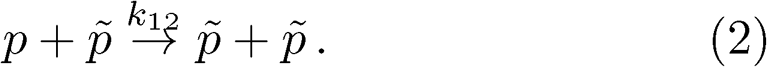

These considerations motivate a system of governing equations for the spatio-temporal evolution of the total amount of healthy and misfolded proteins *p* and 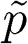,

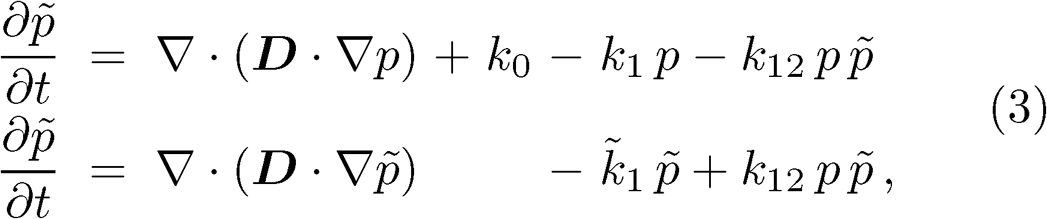

where ***D*** is the diffusion tensor that characterizes protein spreading, *k*_0_ is the production rate of healthy protein, *k*_1_ and 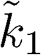 are the clearance rates of *p* and 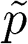, and *k*_12_ is the conversion rate from the healthy to the misfolded state, as shown in Fig. 2. In the initial healthy state, the healthy and misfolded protein concentrations are *p*_0_ = *k*_0_*/k*_1_ and 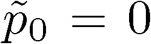; in the diseased state, they converge towards 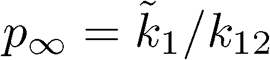 and 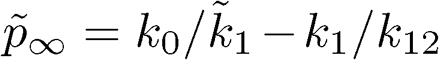. Interestingly, close to the initial healthy state, when 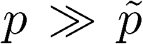, the set of kinetic equations (3) collapses into a single equation of Fisher-Kolmogorov type [15].

**FIG. 2:**
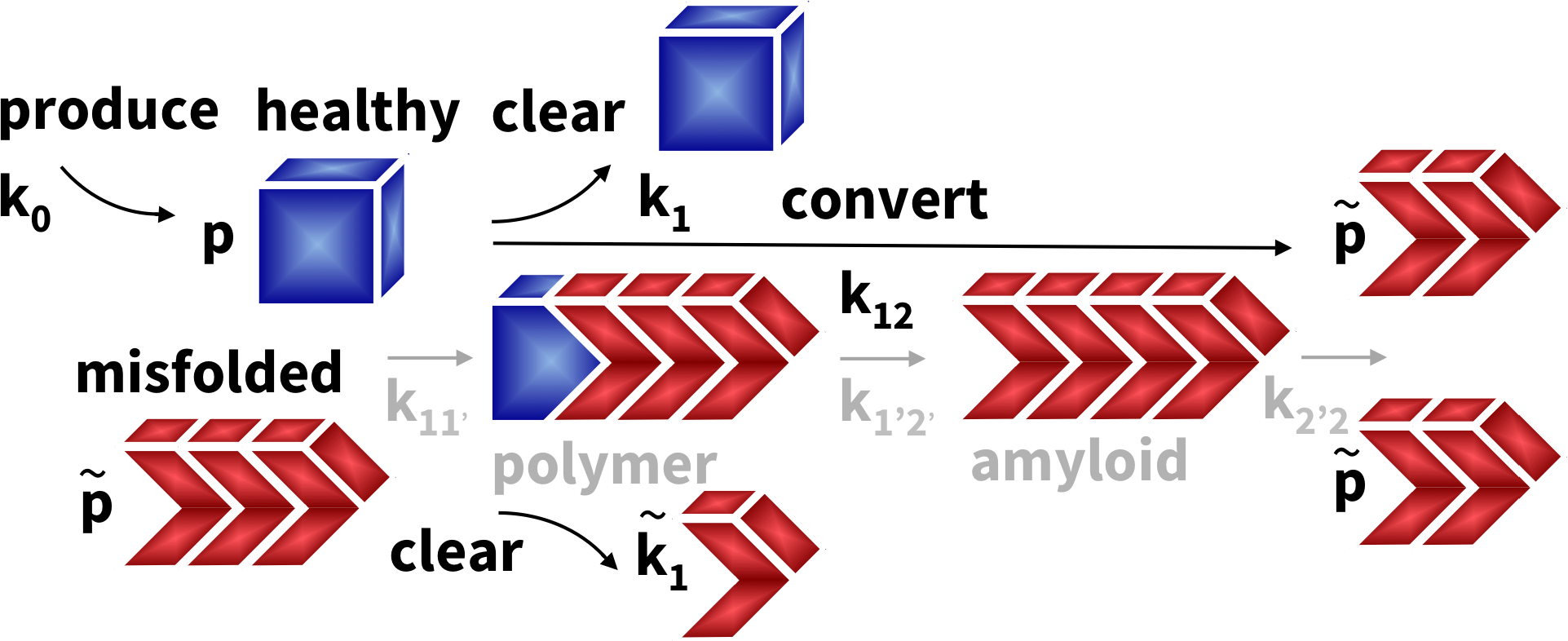
Kinetic model. Misfolded tau proteins 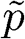 organize in infectious seeds that recruit healthy proteins *p* to misfold on them and then fragment into new seeds.

### Brain network model

We model the spreading of healthy and misfolded proteins as the diffusion across the brain’s connectome, which we represent as a weighted graph 𝒢 with *N* nodes and *E* edges. We extract the graph 𝒢 from the tractography of diffusion tensor images of 418 healthy subjects of the Human Connectome Project [20] using the Budapest Reference Connectome v3.0 [21]. We map the original graph with *N* = 1015 nodes and *E* = 37477 edges onto a graph with *N* = 83 nodes and *E* = 1130 edges in which the degree, the number of edges per node, varies from 6 at the frontal pole to 48 at the caudate. We weight each edge by the mean fiber number *n*_*ij*_ divided by the mean fiber length *l*_*ij*_ averaged over all 418 brains. The mean fiber number varies between 1≤ *n*_*ij*_≤ 596, with an average of 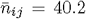 fibers per edge, and most fibers between the superior parietal and precuneus regions. The mean fiber length varies between 11.3 mm ≤*l*_*ij*_≤ 136.8 mm, with an average of 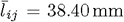. Figure 3 illustrates our graph 𝒢 with the edges color-coded by the mean fiber number *n*_*ij*_, mapped onto a three-dimensional brain model [22].

**FIG. 3:**
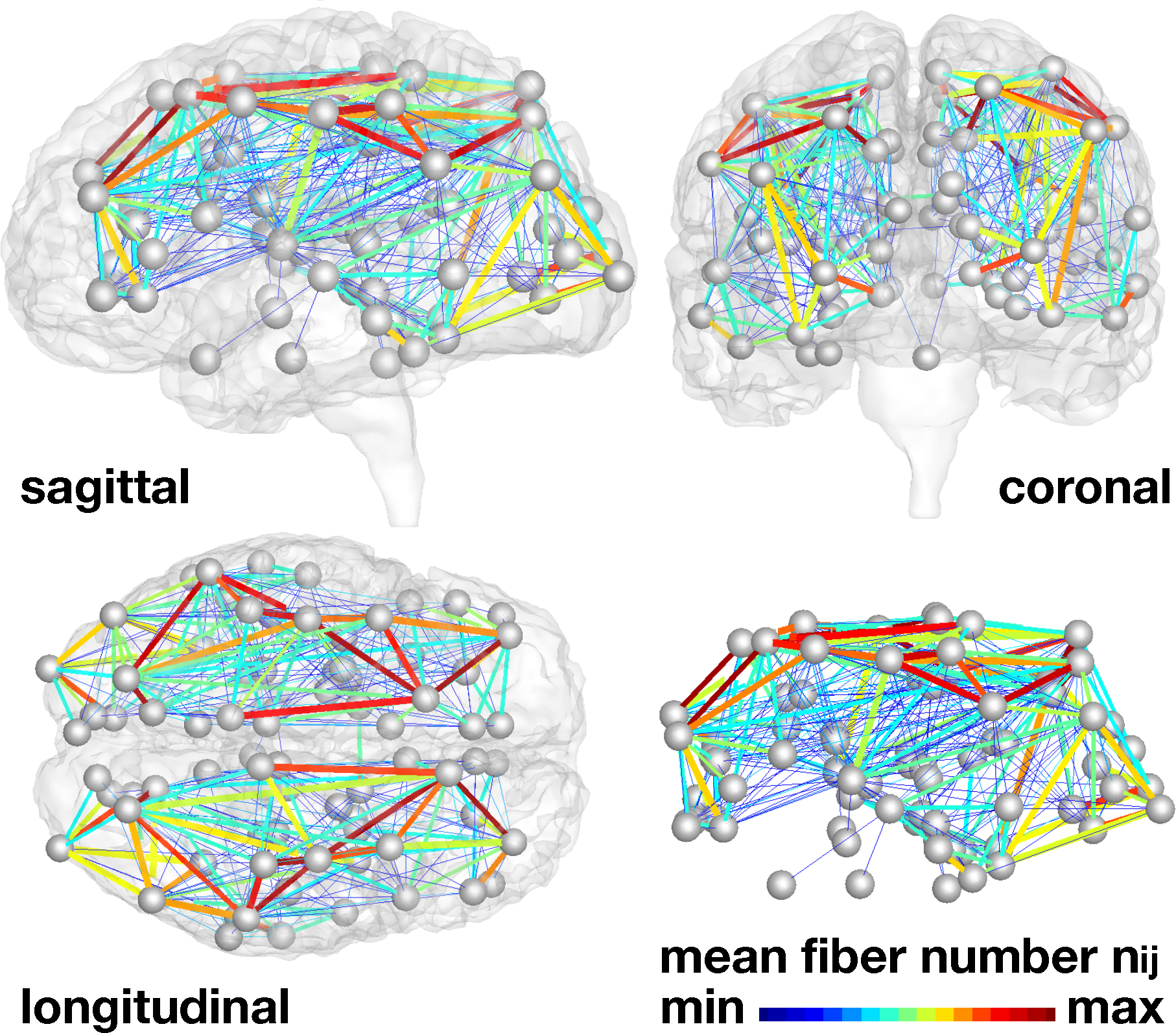
Brain network model. Misfolded tau proteins spread across the brain’s connectome represented as a weighted graph 𝒢 with *N* = 83 nodes and *E* = 1130 edges. Edges are weighted by the mean fiber number *n*_*ij*_ divided by the mean fiber length *l*_*ij*_ averaged over 418 healthy brains.

We can summarize the connectivity of the graph 𝒢 in terms of the degree matrix *D*_*ii*_, a diagonal matrix that characterizes the degree of each node *i*, and the weighted adjacency matrix *A*_*ij*_, the ratio of mean fiber number and length. Their difference defines the weighted graph Laplacian *L*_*ij*_,

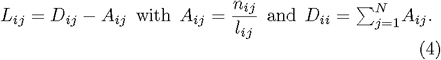

To discretize Eqns. (3) on our weighted graph 𝒢 we introduce the healthy and misfolded protein concentrations *p*_*i*_ and 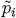 as unknowns at the *i* = 1, *…, N* nodes and assume that the weighted Laplacian *L*_*ij*_ characterizes their spreading across the brain network,

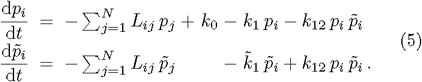

Figure 4 illustrates the degrees *D*_*ii*_ of the non-weighted and connectivity-weighted graphs, left, and the adjacency *A*_*ij*_, right. The degree varies between 2.1 ≤ *D*_*ii*_ ≤ 127.6, with an average degree of 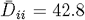 per node, and the lowest and highest degrees in the frontal pole and precentral gyrus, shown in blue and red.

**FIG. 4:**
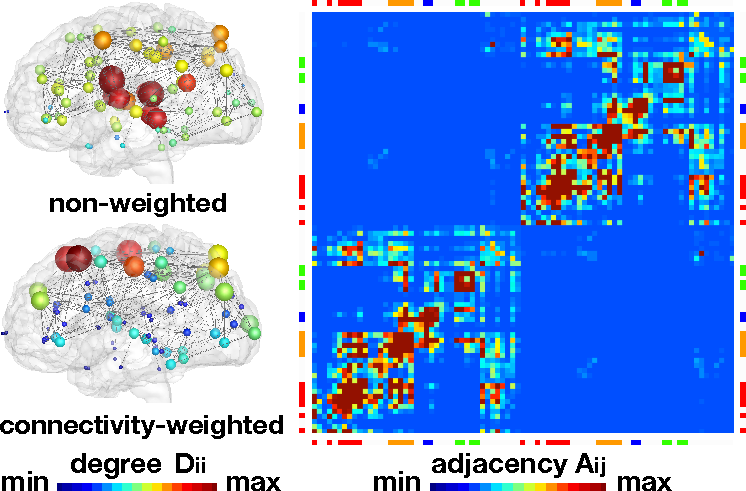
Brain network model. The connectivity of the graph 𝒢 is represented through the degree *D*_*ii*_, the number of edges per node, and the adjacency *A*_*ij*_ = *n*_*ij*_ */l*_*ij*_, the ratio of fiber number and length. Degree *D*_*ii*_ of non-weighted graph (top) and connectivity-weighted graph (bottom), and adjacency *A*_*ij*_ of connectivity-weighted graph (right) averaged over 418 healthy brains.

The adjacency varies between 0.01 ≤ *A*_*ij*_≤ 35.32, with an average adjacency of 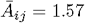 per edge, and lowest and highest values between the superior parietal and precuneus regions and between the lateral orbitofrontal and isthmus cingulate regions. The adjacency matrix clearly reflects the small world architecture of our brain with strongly connected hubs within the right and left hemispheres, the lower left and upper right quadrants, and strong connections within the four lobes, the eight red regions along the diagonal.

### Biomarker model

A biomarker is a global metric to characterize the evolution of neurodegeneration [23], for example, the sum of all misfolded proteins 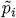 at all *i* nodes,

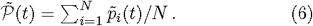

To simulate Alzheimer’s disease, we seed misfolded tau proteins in the entorhinal cortex and allow them to spread across the brain. This takes 30 years in the model and less than a second on a standard laptop computer. The dashed gray and black lines in Fig. 5 show the biomarker of the network model in Fig. 1, bottom, and, for comparison, of the continuum model in Fig. 1, middle. The quantitative comparison of both curves confirms that our network model excellently captures the global characteristics of continuum models for Alzheimer’s disesase [13]. The solid lines in Fig. 5 summarize the biomarker abnormality in all four lobes. The activation sequence, from the temporal to the frontal, parietal, and occipital lobes, agrees well with the neuropathological spreading patterns in Fig. 1, top [11].

**FIG. 5:**
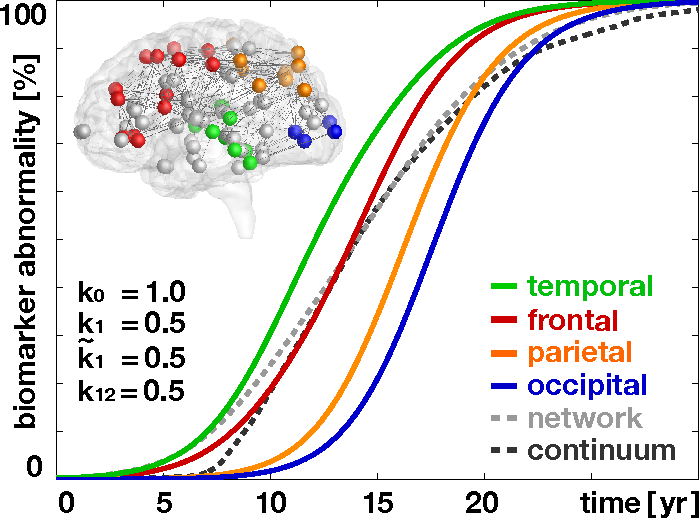
Biomarker abnormality. Summing the concentration of misfolded proteins 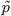 across individual lobes reveals the characteristic activation sequence in Alzheimer’s disease from the temporal lobe to the frontal, parietal, and occipital lobes. The dashed gray and black lines highlight the biomarker abnormality 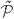 of the network and continuum models in Fig. 1 integrated across the entire brain.

All biomarker curves display a smooth sigmoidal for which is in excellent agreement with clinical biomarke models of neurodegeneration [23].

### Infection times

To characterize the vulnerability of different brain regions, we now seed misfolded proteins in all *N* = 83 regions, simulate their spreading, and calculate their individual biomarker curves and infection times. Fig. 6 summarizes the biomarker curves and their associated brain regions color-coded by infection time. Misfolded proteins spread fastest when seeded in the putamen and insula with a total infection times of 20.2 years, shown in red, and slowest when seeded in the frontal pole and entorhinal region with infection times of 30.4 and 28.8 years, shown in blue. This significant variation in infection time underlines the heterogeneity of our brain network model [24]. For comparison, the dashed gray line illustrates the lower limit of the infection time of 16.6 years, associated with a homogeneous seeding across all *N* = 83 regions. The entorhinal cortex, the region where misfolded tau proteins are first observed [11], is associated with the second longest in infection time. This could explain–at least in part–why tau pathology is so slow and difficult to diagnose during the early stages of Alzheimer’s disease [8].

**FIG. 6:**
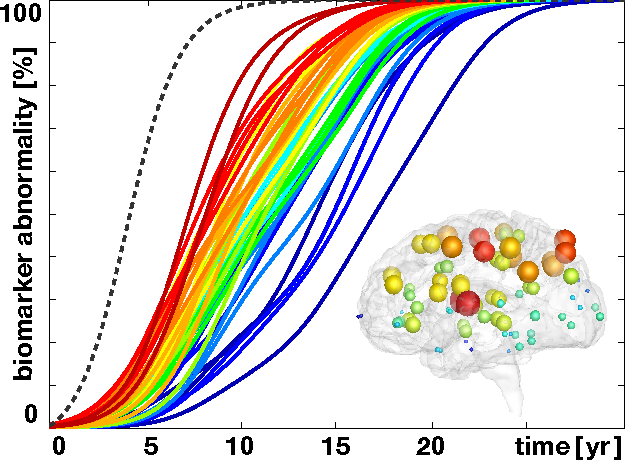
Infection time. Biomarker abnormality for *N* = 83 seeding regions illustrates the regional vulnerability of the brain network. The dashed gray line highlights the lower limit of the infection time associated with a homogeneous seeding across all regions.

### Treatment opportunities

Two promising therapeutic strategies are currently emerging to delay or even prevent the progression of Alzheimer’s disease [25]: reducing misfolding [26] and increasing clearance [27]. Figures 7 and 8 probe the effect of reduced misfolding. A turnover rate of *k*_12_ = 0.50 predicts the baseline progression of Alzheimer’s disease in agreement with Fig. 1. Decreasing the turnover rate to *k*_12_ = 0.45 and *k*_12_ = 0.40 delays and reduces the accumulation of misfolded tau protein 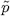 and with it the biomarker abnormalit 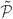. In the early stages of neurodegeneration, even a small reduction of misfolding can delay disease progression by several decades [26] and reduce the resting state of misfolded protein 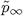 to 89% and 75% of its baseline value of Alzheimer’s disease. Figures 9 and 10 probe the effect of increasing the clearance of misfolded tau protein 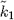. A clearance rate of 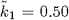 predicts the baseline progression of Alzheimer’s in agreement with Fig. 1. Increasing the clearance rate to 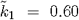 and 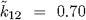 has similar effects as decreasing the turnover rate *k*_12_; it delays and reduces the accumulation of misfolded tau protein 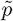 and with it the biomarker abnormality 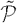, see Supplemental Movies. Similar to a decreased turnover, an increased clearance can delay disease progression by several decades [27] and reduce the resting state of misfolded protein 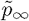 to 67% and 43% of its baseline value of Alzheimer’s disease. It would be interesting to explore how our simulations would change if we not only modeled tau but the combined effects of tau and amyloid beta aggregation [12, 25].

**FIG. 7:**
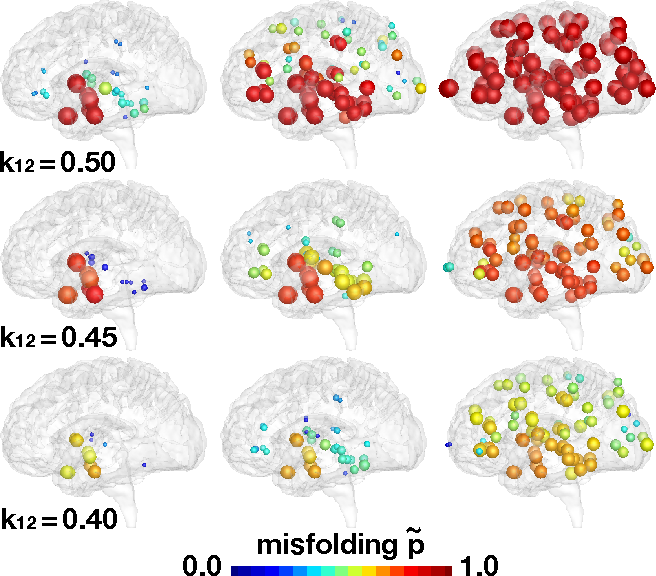
Reducing misfolding. Lower turnover rates *k*_12_ delay and reduce the accsumulation of misfolded tau protein 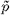. Baseline Alzheimer’s disease (top) and Alzheimer’s disease with moderately (middle) and markedly (bottom) reduced turnover *k*_12_ from healthy to misfolded tau protein.

**FIG. 8:**
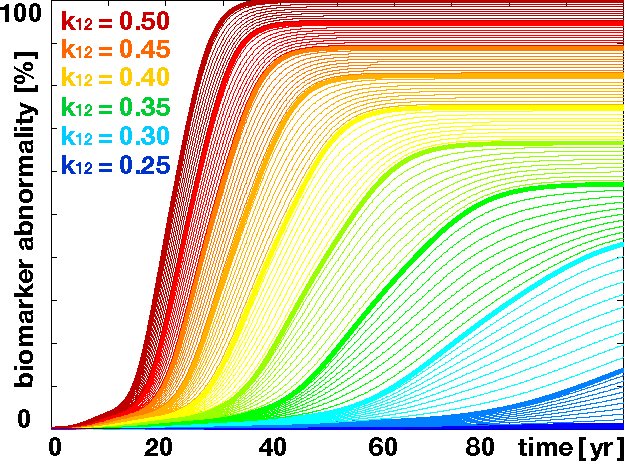
Reducing biomarker abnormality through reduced misfolding. Decreasing the turnover *k*_12_ delays and reduces the accumulation of misfolded tau protein 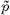 and with it the biomarker abnormality 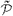.

**FIG. 9:**
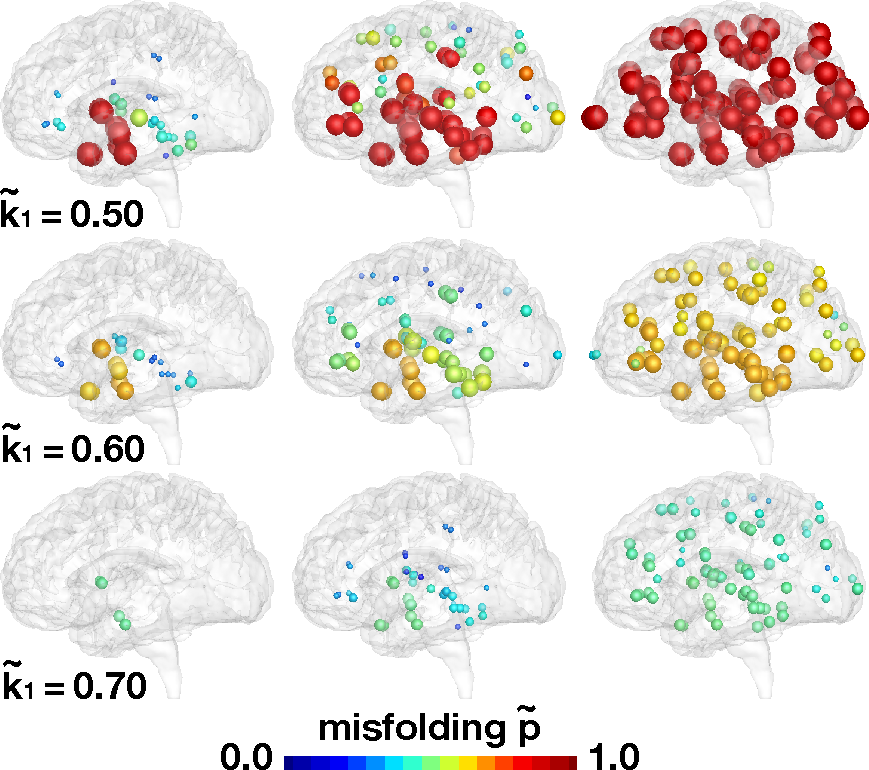
Increasing clearance. Higher clearance rates 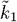 delay and reduce the accumulation of misfolded tau protein 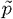. Baseline Alzheimer’s disease (top) and Alzheimer’s disease with moderately (middle) and markedly (bottom) increased clearance 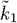 of misfolded tau protein.

**FIG. 10:**
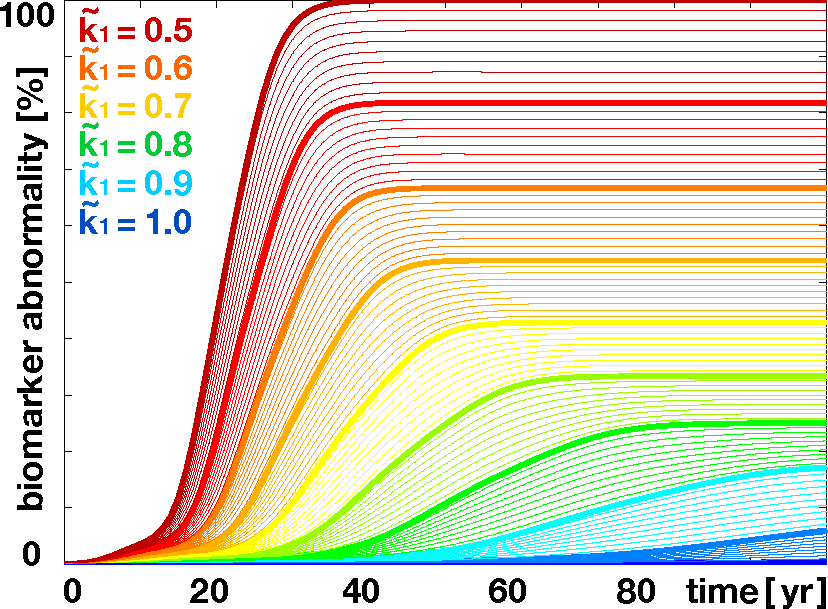
Reducing biomarker abnormality through in-creased clearance. Increasing the clearnace 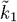 delays and reduces the accumulation of misfolded tau protein 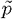 and with it the biomarker abnormality 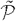.

## Conclusions

Despite their complexity, neurodegenerative diseases display remarkably consistent damage and atrophy patterns. In Alzheimer’s disease, these invasion patterns are highly correlated with the spreading pattern of misfolded tau proteins. Here we model the spreading of tau proteins by combining misfolding kinetics and network diffusion through a connectivity-weighted graph. In our dynamic brain network model, the concentrations of healthy and misfolded proteins emerge dynamically at each node and propagate across the graph through its connectivity-weighted edges. Our model correctly predicts the spatio-temporal spreading pattern of tau in Alzheimer’s disease. Its computational efficiency allows us to rapidly screen the landscape of process parameters that govern the kinetics of protein misfolding. We demonstrate the potential of our model by simulating biomarker curves, infection times, and therapeutic intervention. A better understanding of the spreading of misfolded proteins could open new therapeutic opportunities towards blocking protein misfolding and promoting protein clearance using antibodies or small molecules.

## Acknowledgments

This work was supported by the Engineering and Physical Sciences Research Council grant EP/R020205/1 to Alain Goriely and by the National Science Foundation grant CMMI 1727268 to Ellen Kuhl.

## References

[1] D. S. Bassett and E. Bullmore, The Neuroscientist 12, 512 (2006).

[2] S. Bassett and E. T. Bullmore, The Neuroscientist 23, 499 (2017).

[3] Bullmore and O. Sporns, Nature Reviews Neuroscience 10, 186 (2009).

[4] A. Raj, A. Kuceyeski, and M. Weiner, Neuron 73, 1204 (2012).

[5] Y. Iturria-Medina, R. C. Sotero, P. J. Toussaint, A. C. Evans, and A. D. N. Initiative, PLoS Computational Biology 10, e1003956 (2014).

[6] M. Jucker and L. C. Walker, Annals of Neurology 70, 532 (2011).

[7] S. B. Prusiner, Proceedings of the National Academy of Sciences 95, 13363 (1998).

[8] L. C. Walker and M. Jucker, Annual Review of Neuroscience 38, 87 (2015).

[9] M. Goedert, Science 349, 1255555 (2015).

[10] P. C. Bressloff and J. M. Newby, Reviews of Modern Physics 85, 135 (2013).

[11] H. Braak and E. Braak, Acta Neuropathologica 82, 239 (1991).

[12] M. Jucker and L. C. Walker, Nature 501, 45 (2013).

[13] J. Weickenmeier, E. Kuhl, and A. Goriely, Physical Review Letters 121,158101 (2018).

[14] T. P. Knowles, C. A. Waudby, G. L. Devlin, S. I. Cohen, A. Aguzzi, M. Vendruscolo, E. M. Terentjev, M. E. Welland, and C. M. Dobson, Science 326, 1533 (2009).

[15] J. Weickenmeier, M. Jucker, A. Goriely, and E. Kuhl, Journal of the Mechanics and Physics of Solids 124, 264 (2019).

[16] R. A. Fisher, Annals of Eugenics 7, 355 (1937).

[17] A. N. Kolmogorov, Moscow University Bulletin of Mathematics 1, 1 (1937).

[18] F. Matthäus, Journal of Theoretical Biology 240, 104 (2006).

[19] P. C. Bressloff, Lecture Notes on Mathematical Modelling in the Life Sciences (2014).

[20] J. A. McNab, B. L. Edlow, T. Witzel, S. Huang, H. Bhat, K. Heberlein, T. Feiweier, K. Liu, B. Keil, J. Cohen-Adad, et al., Neuroimage 80 (2013).

[21] B. Szalkai, C. Kerepesi, B. Varga, and V. Grolmusz, Cognitive Neurodynamics 11, 113 (2017).

[22] J. Weickenmeier, C. Butler, P. G. Young, A. Goriely, and E. Kuhl, Computer Methods in Applied Mechanics and Engineering 314, 180 (2017).

[23] C. R. Jack and D. M. Holtzman, Neuron 80, 1347 (2013).

[24] M. Barthelemy, A. Barrat, R. Pastor-Satorras, and Vespignani, Physical Review Letters 92, 178701 (2004).

[25] J. C. Polanco, L. G. Bodea, R. Martinez-Marmol, F. A. Meunier, and J. Götz, Nature Reviews Neurology 14, 22 (2018).

[26] E. E. Congdon and E. M. Sigurdsson, Nature Reviews Neurology 14, 399 (2018).

[27] S.-H. Xin, L. Tan, X. Cao, J.-T. Yu, and L. Tan, Cognitive Neurodynamics 34, 733 (2018).

